# IPPK-1 and IP6 contribute to ventral nerve cord assembly in *C. elegans*

**DOI:** 10.1101/2024.03.03.583222

**Authors:** Nathaniel Noblett, Tony Roenspies, Stephane Flibotte, Antonio Colavita

**Affiliations:** Neuroscience Program, Ottawa Hospital Research Institute, Ottawa, Canada; Department of Cellular and Molecular Medicine, University of Ottawa, Ottawa, Canada; University of Ottawa Brain and Mind Research Institute, Ottawa, Canada; UBC/LSI Bioinformatics Facility, University of British Columbia, Vancouver, Canada

**Keywords:** Convergent extension, nerve cord, inositol phosphate, IPPK, IP6

## Abstract

Inositol phosphates (IPs) are essential for the development and function of the nervous system. Loss-of-function studies, which demonstrate the importance of specific IP isomers, show their critical role in proper neural tube formation. In this study, we show that inositol pentakisphosphate 2-kinase (IPPK-1), the kinase that phosphorylates IP5 to generate IP6, is involved in assembling the ventral nerve cord (VNC) in *C. elegans*. We show that mutations in *ippk-1* lead to the mispositioning of motor neurons along the VNC of newly hatched larvae. These positioning defects reflect disruption of VNC assembly during embryogenesis, as VNC neuronal progenitors in *ippk-1* embryos display a more compact organization after arising on the left and right sides of the embryo, delays in rosette-mediated convergent extension, and defects in cell intercalation. We further show that injection of exogenous IP6 into the gonads of *ippk-1* mutants can rescue both embryonic and neuron positioning defects. Our findings indicate that IP isomers, particularly IP6, are important for ventral nerve cord formation in *C. elegans*. Along with their role in neural tube formation in vertebrates, these results suggests that IP isomers play an ancient role in central nerve cord development.

**Highlights:**

- *ipmk-1* and *ippk-1* mutants display neuron position defects in the ventral nerve cord (VNC).
- *ippk-1* mutants display disorganization in VNC neuronal progenitors during VNC assembly.
- IPPK-1 is involved in convergent extension during VNC formation.
- Exogenous IP6 rescues larval and embryonic defects in *ippk-1* mutants.

## Introduction

Inositol phosphates and polyphosphates, groups of metabolites produced through the progressive phosphorylation of inositol triphosphate (IP3), are central players in a diverse range of cellular processes. These include the regulation of ion channel activity, cell migration, exocytosis, developmental timing and the establishment of left-right asymmetry (Efanov et al., 1997; Yang et al., 2001; Sarmah et al., 2005; Jadav et al., 2016; Yang et al., 2021; Rao et al., 2015). These functions are essential for proper embryogenesis and organogenesis during development (Yang et al., 2021; Frederick et al., 2005; Boitano et al., 1992; Seeds et al., 2015). In the developing nervous system, the activity of inositol phosphate and polyphosphate kinases regulate several aspects of brain development and function (Loss et al., 2013; Park et al., 2019a; Park et al., 2019b; Ahmed et al., 2015; Wilson et al., 2009). Regulation of inositol phosphate levels, specifically inositol pentakisphosphate (IP5) and inositol hexaphosphate (IP6), are important for neuronal survival and differentiation (Loss et al., 2013; Ucuncu et al., 2020). Additionally, loss of inositol hexaphosphate kinases lead to impaired brain development due to neuronal migration, morphology and synapse formation defects (Rojas et al., 2019; Fu et al., 2017; Fu et al., 2015).

Neural tube defects, caused by a failure of the neural plate to fold, fuse or undergo canalization during neurulation, are highly prevalent congenital abnormalities (reviewed in Greene et al., 2017; Nikolopoulou et al., 2017). Disrupting inositol phosphate metabolism, either through deprivation of dietary inositol or introducing loss-of-function mutations in IP synthesis enzymes, results in or exacerbates cranial neural tube defects (NTDs) (Wilson et al., 2009; Cockroft et al., 1992; Greene and Copp, 1997). Several studies have also described functions for highly phosphorylated forms of inositol during neural tube closure. Loss of inositol-tetrakisphosphate 1-kinase (ITPK1) and inositol polyphosphate multikinase (IPMK), kinases required for the production of IP5 and its derivatives, results in lethality around the neurulation stage in mice and has been linked to an increase in NTD cases in humans (Wilson et al., 2009, Frederik et al., 2005, Guan et al., 2014; Verbsky et al., 2005a).

While numerous genes have been implicated in the development of NTDs in vertebrates, a key category are those that encode for proteins involved in convergent extension. This morphogenetic process is marked by the constriction of cell-cell junctions and cell intercalation that act to narrow a tissue along one axis and lengthen it along another (reviewed in Sutherland et al., 2020). Coordination of convergent extension across the neural tube is a prominent driver of early neurulation, which depends on the closure of neuroepithelial folds. Failure of this process can lead to severe NTDs in vertebrates (Wallingford and Harland, 2002; Goto and Keller, 2002). Components of the conserved Wnt-planar cell polarity (PCP) pathway, such as VANGL1/2 and their orthologues, play important roles in coordinating convergent extension in mice (Lei et al., 2019; Curtin et al., 2003; Murdoch, 2003), *Xenopus* (Wallingford and Harland, 2002; Williams et al., 2014; Butler and Wallingford, 2018) and zebrafish (Ciruna et al., 2006; Reynolds et al., 2010). Besides PCP signaling, junctional contraction during convergent extension are also controlled by transient Ca^2+^ oscillations resulting from the interaction between inositol triphosphate (IP3) and its receptor (Cogram et al., 2004; Politi et al., 2006, Mound et at al., 2017). In support of this, reducing Ca2+ waves through pharmacological or genetic means impairs axial elongation and disrupts convergent extension (Wallingford et al., 2001; Westfall et al., 2003). However, the role that more highly phosphorylated forms of inositol play during neural tube development remains poorly understood.

In *C. elegans*, convergent extension movements are involved in the early assembly of the ventral nerve cord (VNC) (Shah et al., 2017). The VNC runs along the anterior-posterior (AP) axis of the worm and is comprised of three classes of motor neurons (DD, DA and DB) at hatching. During embryogenesis, neuronal progenitors born on the left and right sides of the embryo, undergo cell-cell intercalations, including processes driven by the formation and resolution of multicellular rosettes, to converge toward and extend along the midline. Following this, progenitor cell bodies are positioned along the AP axis in a predominantly single-file formation. This process involves VANG-1/PCP and SAX-3/Robo-mediated regulation of cell-cell intercalations (Shah et al., 2017). Loss of VANG-1 or SAX-3 signaling delays rosette resolution and, subsequently, the single-file intercalation of progenitors at the midline. However, simultaneous disruption of both pathways leads to a more severe convergent extension defect, which, in the most extreme cases, results in the anterior displacement of most motor neuron cell bodies in the VNC at hatching (Shah et al., 2017).

In this study, we explore the role of *ippk-1*, the gene encoding the worm orthologue of mammalian inositol-pentakisphosphate 2-kinase (IPPK/IP5K), in VNC assembly (Davis et al., 2022). IPPK is an enzyme in the IP biosynthesis pathway responsible for phosphorylating IP5 to generate IP6 (Verbsky et al., 2002). Our study reveals that the loss of *ippk-1* results in the mispositioning of motor neuron cell bodies within the VNC of newly hatched worms. Disruption of the worm orthologue of IPMK also produces a similar phenotype. Examination of VNC morphogenesis in *ippk-1* mutant embryos uncovered a more compact organization as VNC neuronal progenitors from the left and right sides migrated toward the midline. Additionally, we observed defects related to abnormal convergent extension, including delays in rosette resolution, persistent cell contacts after midline intercalation, and reduced VNC extension by the 1.5-fold stage. Motor neuron positioning defects in newly hatched worms, a consequence of abnormal VNC development during embryogenesis, were rescued by the addition of exogenous IP6. These findings highlight the importance of IP biosynthesis pathways and polyphosphorylated IPs in the development of central nerve cords.

## Results

### IPPK-1 is required for proper positioning of DD neurons along the VNC

At hatching, the VNC contains three classes of motor neurons (6 GABAergic inhibitory DD, 9 cholinergic excitatory DA, and 7 cholinergic excitatory DB) arranged in repeating DD-DA-DB motor pools responsible for sinusoidal locomotion (Lu et al., 2022). The cell bodies of these neurons are stereotypically positioned along the VNC (Saharkhiz et al., 2024). We have previously shown that loss of planar cell polarity genes like *vang-1*/VANG result in VNC assembly defects during embryogenesis which manifest as mispositioned motor neurons in L1 larvae (Shah et al., 2017). To identify new genes involved in neuron positioning, we performed a genetic screen (A. Colavita, unpublished results) using the DD-specific reporter *ynIs37[flp-13p::GFP]* (a gift from Dr. Chris Li, CCNY) and identified *zy65*. Whole genome sequencing revealed *zy65* to be mutation in *ippk-1*, the sole *C. elegans* orthologue of inositol-pentakisphosphate 2-kinase (IPPK), a kinase in the IP biosynthesis pathway that catalyzes the conversion of IP5 to IP6 (Fig. 1A and B)*. ippk-1(zy65)* contains an A to T nucleotide change in the consensus splice acceptor site of exon 6 that is predicted to affect splicing of the *ippk-1* transcript (Fig. 1A).

**Figure 1.**
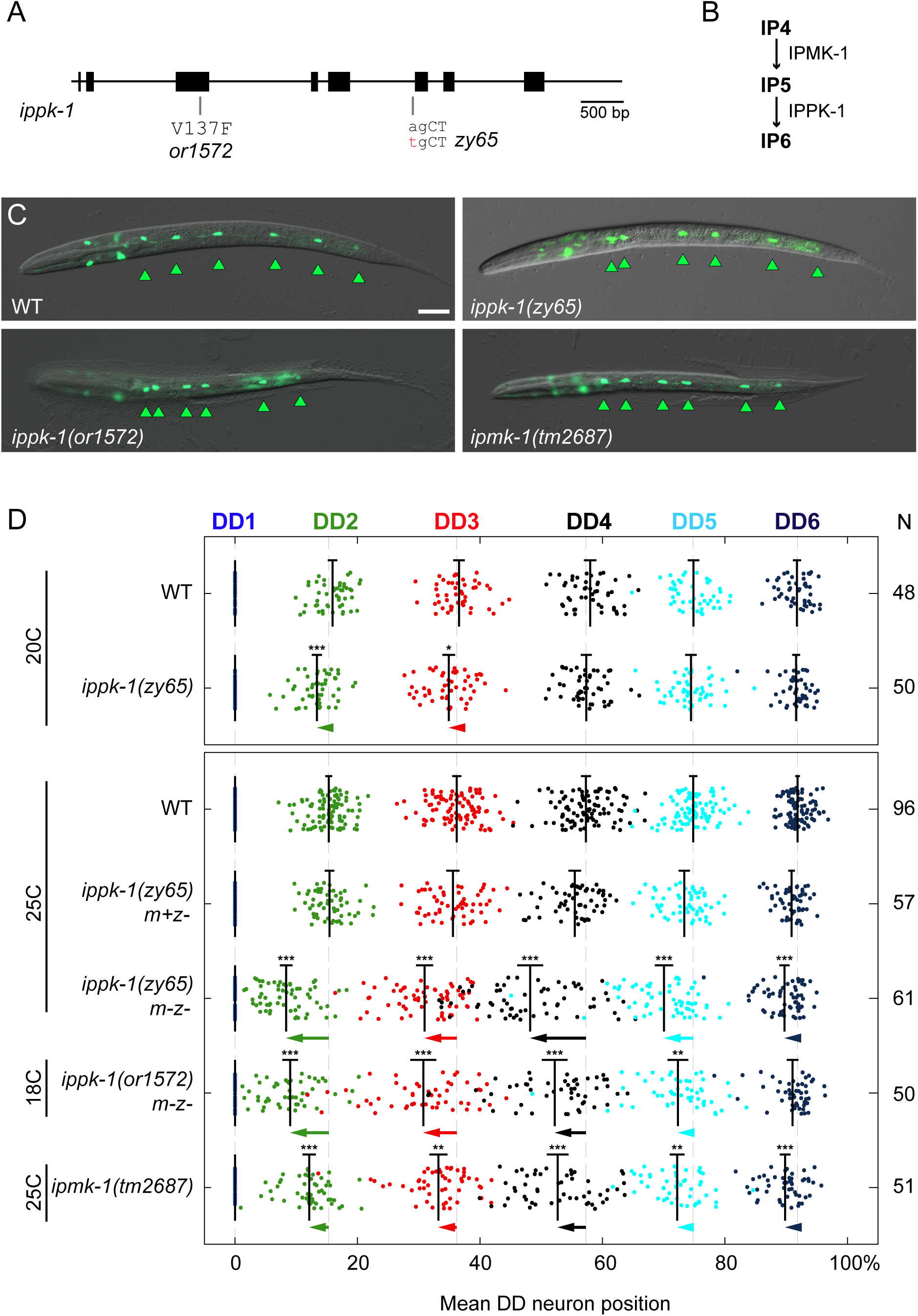
*ippk-1* mutants display DD neuron spacing defects. (A) Diagram of the *ippk-1* genomic region labeled with alleles used in this study. (B) Kinase cascade pathway involved in phosphorylating IP4 and IP5 to produce IP6. (C) Representative images of DD positions in WT, *ippk-1* and *ipmk-1* mutants, visualised using the DD-specific *flp-13p::GFP* reporter. Arrowheads mark DD neurons. Scale bar = 20μm. (D) Quantification of DD2–DD6 mean positions relative to DD1 in L1 stage WT and mutant worms. Neurons (colour coded as indicated along top) are plotted along the AP axis, where DD1 and anus mark the 0% and 100% positions respectively. Means and 95% confidence intervals are indicated for each neuron. Animals scored (N) indicated on the right. Statistics: One-way ANOVA with Dunnett’s post-hoc test against the corresponding WT neuron, except for strains scored at 20⁰C, where a two-tailed t-test with Welch’s correction was used. *p<0.05, **p<0.01, ***p<0.001. Arrows indicate the size of the shift from the WT mean for all neurons where p>0.05.

IPPK-1 was previously shown to be essential for larval growth and in the maintenance of adult germline membrane architecture (Lowry et al., 2015). Our findings indicate an additional role in VNC development. In newly hatched animals, DD motor neurons display an approximately equidistant spacing along the VNC. In *zy65* mutants, DD neurons are located at more anterior positions compared to wild type (WT) animals (Fig. 1C and D). This phenotype is temperature sensitive as there is a significant increase in the penetrance of anteriorly shifted DD neurons when animals are incubated at 25°C compared to 20°C (Fig. 1D). We also found that DD position defects are maternally rescued indicating a role for maternally derived IPPK-1 or its product IP6 (Fig. 1D). To determine if the changes in neuron position are unique to DD neurons or shared by other motor neuron classes, we also examined DB neurons using an mScarlet knock-in at the *vab-7* locus (Saharkhiz et al., 2024) (Fig. S1). DB neurons were also mispositioned more anteriorly in *zy65* mutants, indicating a more general role in motor neuron positioning.

Two observations confirm that the *ippk-1* gene is responsible for the observed motor neuron position defects. First, the position defects in *ippk-1(zy65)* were rescued by expressing the *ippk-1* cDNA from either an *ippk-1* or pan-neuronal *unc-33* promoter (Fig. S2). Second, *ippk-1(or1572)*, a temperature-sensitive allele from the Lowry et al. (2015) study, also displayed similar anterior shifts in DD cell body positions (Fig. 1C and D). As *or1572* exhibited sterility and larval arrest at temperatures above 20°C, it was scored at 18°C to ensure the production of viable larvae. *ippk-1(zy65*) worms also displayed sterility and epidermal morphology defects, similar to those in *ippk-1(or1572)*, when grown at the restrictive temperature. 9.5% (N=116) of post-comma stage embryos and 17.4% (N=135) of larvae displayed epidermal morphology defects characterized by a lumpy appearance, with lumps mostly in the posterior (Fig. S3). Notably, loss of upstream components required for the generation of inositol triphosphate (IP3) have been reported to show similar defects in both embryonic and larval stages (Vázquez-Manrique et al., 2008).

Since IPPK-1 acts in an IP synthesis pathway with several other kinases, we also asked if loss of another kinase in this pathway would also disrupt DD positioning. IPMK phosphorylates both IP3 and IP4 to produce IP5 (Saiardi et al., 1999; Odom et al., 2000) (Fig. 1B). *ipmk-1* is the only *C. elegans* orthologue of IPMK. Unlike *ippk-1*, which is an essential gene, the *ipmk-1(tm2687)* deletion allele (Mitani, 2017), a probable null, is viable (Yang et al., 2021). We found that *tm2687* displayed DD position defects similar to those in *ippk-1* mutants (Fig. 1C and D). As human IPPK has been shown to play additional non-catalytic roles unrelated to IP synthesis (Brehm et al., 2013), these findings indicate that an intact IP kinase cascade is important for proper DD positioning in the VNC.

### *ippk-1* is expressed at low levels, except in the spermatheca

To determine where *ippk-1* is expressed, we generated a transcriptional reporter and tagged the endogenous *ippk-1* locus with GFP (Fig. S4). The transcriptional reporter was made using approximately 3kb of *ippk-1* promoter fused to GFP (Fig. S4A). Expression from a transgenic array was observed in pharynx, intestine, spermatheca, and a subset of head, tail and ventral cord neurons (Fig. S4B-D). Two GFP knock-in strains were made, with GFP inserted at either the N or C-terminus of *ippk-1*, using the CRISPR/Cas9 homology-directed repair approach described in Dickinson et al. (2015). This approach utilizes a selection cassette (SEC) containing a GFP reporter, followed by a loxP-flanked stop cassette which allows transcriptional read-through past the reporter only upon Cre recombinase-mediated SEC excision. The N-terminal insertion can therefore function as a transcriptional reporter prior to SEC excision. In contrast to the transcriptional reporter, the *ippk-1(zy100[GFP::loxP-stop-loxP::IPPK-1])* strain, which reflects endogenous promoter activity, showed strong expression only in the spermatheca (Fig. S4E). Likewise, both the N-terminal (*zy101*) and C-terminal (*zy102*) GFP knock-ins appeared to show protein expression only in the spermatheca (Fig. S4F). With the exception of the spermatheca, we did not observe expression in other areas of the somatic gonad or germline, where we might have expected it, given its role in germline architecture and gonad arm morphology (Lowry et al., 2015). This observation is largely consistent with single-cell transcriptome data, which shows a generally low abundance of *ippk-1* transcripts except in the spermatheca (Taylor et al., 2021; Packer et al., 2019).

### IPPK-1 acts in concert with VANG-1 and SAX-3 to position DD neurons in the VNC

We previously demonstrated that the PCP pathway protein VANG-1 and the Robo receptor SAX-3 function in parallel to ensure proper motor neuron cell body positioning in the VNC of newly hatched worms. Simultaneous deletion of both *vang-1* and *sax-3* leads to the highly penetrant displacement of motor neuron cell bodies toward the anterior, which is more severe than the defect caused by loss of either gene alone (Fig. 2) (Shah et al., 2017). This striking anterior displacement is the result of a severe disruption of convergent extension movements as neuronal progenitors undergo mediolateral convergence and anterior-posterior extension to form the VNC during embryogenesis (Shah et al., 2017). To investigate genetic interactions between *ippk-1* and these pathways, we examined DD neuron positions in *ippk-1* double mutants with *vang-1* and *sax-3*. We found that double mutants containing *ippk-1(zy65)* displayed significantly stronger anterior shifts, at one or more DD neuron cell body positions, compared to *vang-1* and *sax-3* single mutants (Fig. 2). A similar, albeit milder, increase in severity was also observed in *vang-1; ipmk-1* and *sax-3; ipmk-1* double mutants (Fig. 2). Interestingly, although *vang-1* and *sax-3* double mutants containing *ippk-1(zy65)* displayed similar shifts in DD positions, double mutants containing *ippk-1(or1572)* displayed stronger shifts in the *vang-1* mutant background than in the *sax-3* background. This difference appeared to be absent in *ipmk-1* double mutants, suggesting that the *zy65* and *or1572* lesions disrupt IPPK-1 protein function in different ways, with *or1572* likely retaining more kinase activity. Overall, these results are consistent with an *ipmk-1* and *ippk-1* containing IP pathway acting, at least in part, in parallel with *vang-1* and *sax-3* to promote proper motor neuron positioning in the VNC.

**Figure 2.**
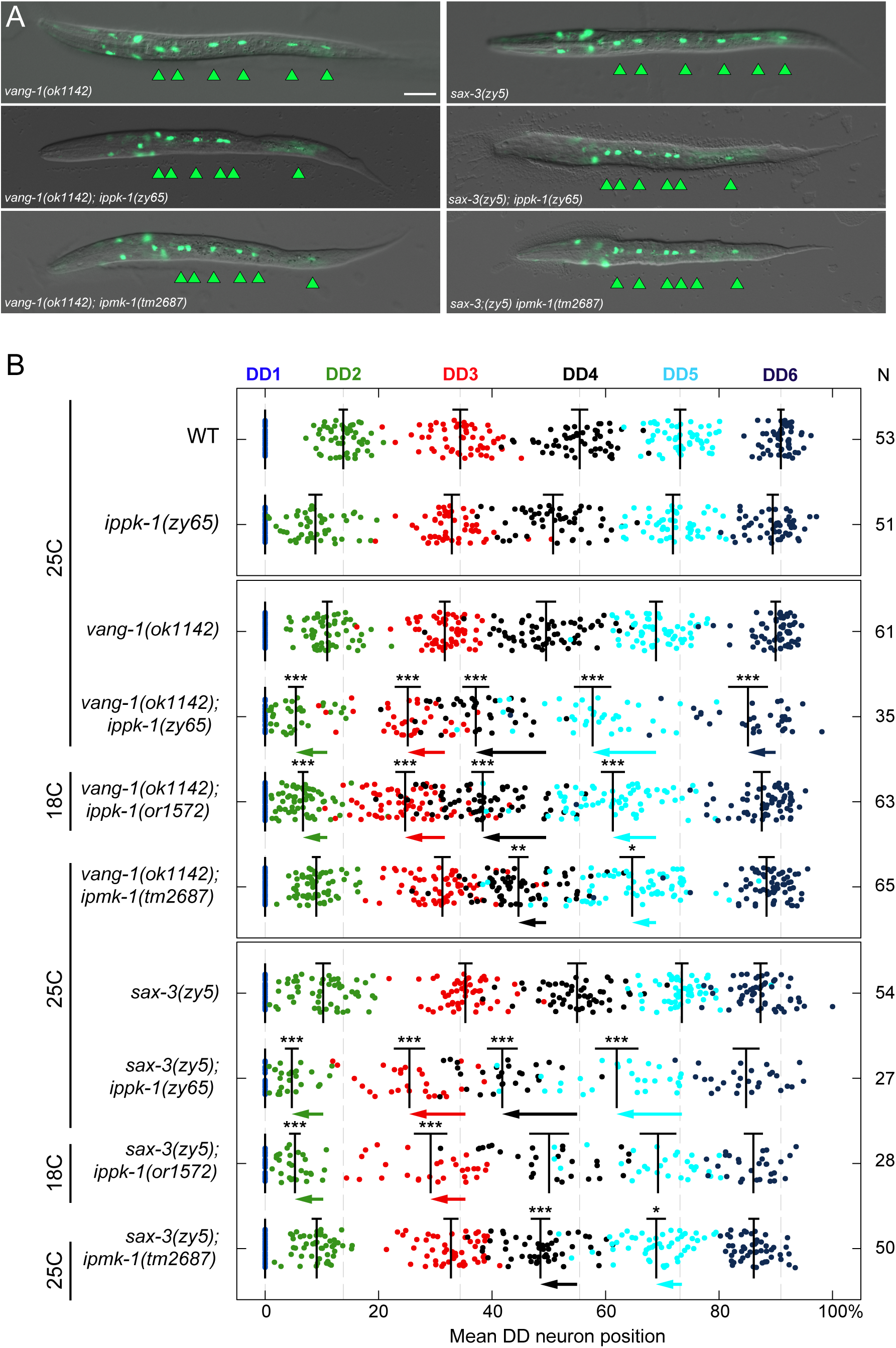
DD position defects are more severe in *vang-1; ippk-1* and *sax-3; ippk-1* double mutants compared to single mutants. (A) Representative images and (B) quantification of mean DD neuron position relative to DD1 in *vang-1* and *sax-3* single mutants or in combination with *ippk-1* and *ipmk-1* mutants. Scale bar = 20μm. Means, 95% confidence intervals and plot format as in Fig. 1D. Statistics: One-way ANOVA with Tukey’s post-hoc test. The significance of each neuron across all double mutants is shown relative to the position of its constituent *vang-1(ok1142)* or *sax-3(zy5)* single mutant. *p<0.05, **p<0.01, ***p<0.001. Arrows indicate the size of the mean position shift in double mutants compared to the relevant *vang-1* or *sax-3* single mutant where p>0.05.

### *ippk-1* mutants display defective organization of DD and DA progenitors at midline contact

During embryogenesis, DD and DA progenitors from both the left and right sides, form tightly juxtaposed groups that migrate towards the midline, where they intercalate to establish the presumptive VNC (Shah et al., 2017). To begin to understand how *ippk-1* regulates VNC formation during this stage, we performed time-lapse microscopy using a *cnd-1p::PH::mCherry* reporter to label the membranes of DD and DA progenitors (specifically, DD1-6 and DA1-5) (Fig. 3A). In *ippk-1* mutants, we found that these progenitors appeared more disorganized and compact as they moved toward the midline (Fig. 3B). To quantify tissue compactness, we measured the combined area occupied by the progenitors on the left and right sides in each embryo at the time of midline contact. We found that in *ippk-1(zy65)* mutants, this area was significantly smaller (119.1 µm^2^) (N=12, p<0.0001) compared to WT (151.8 µm^2^) (N=13) (Fig. 3C).

**Figure 3.**
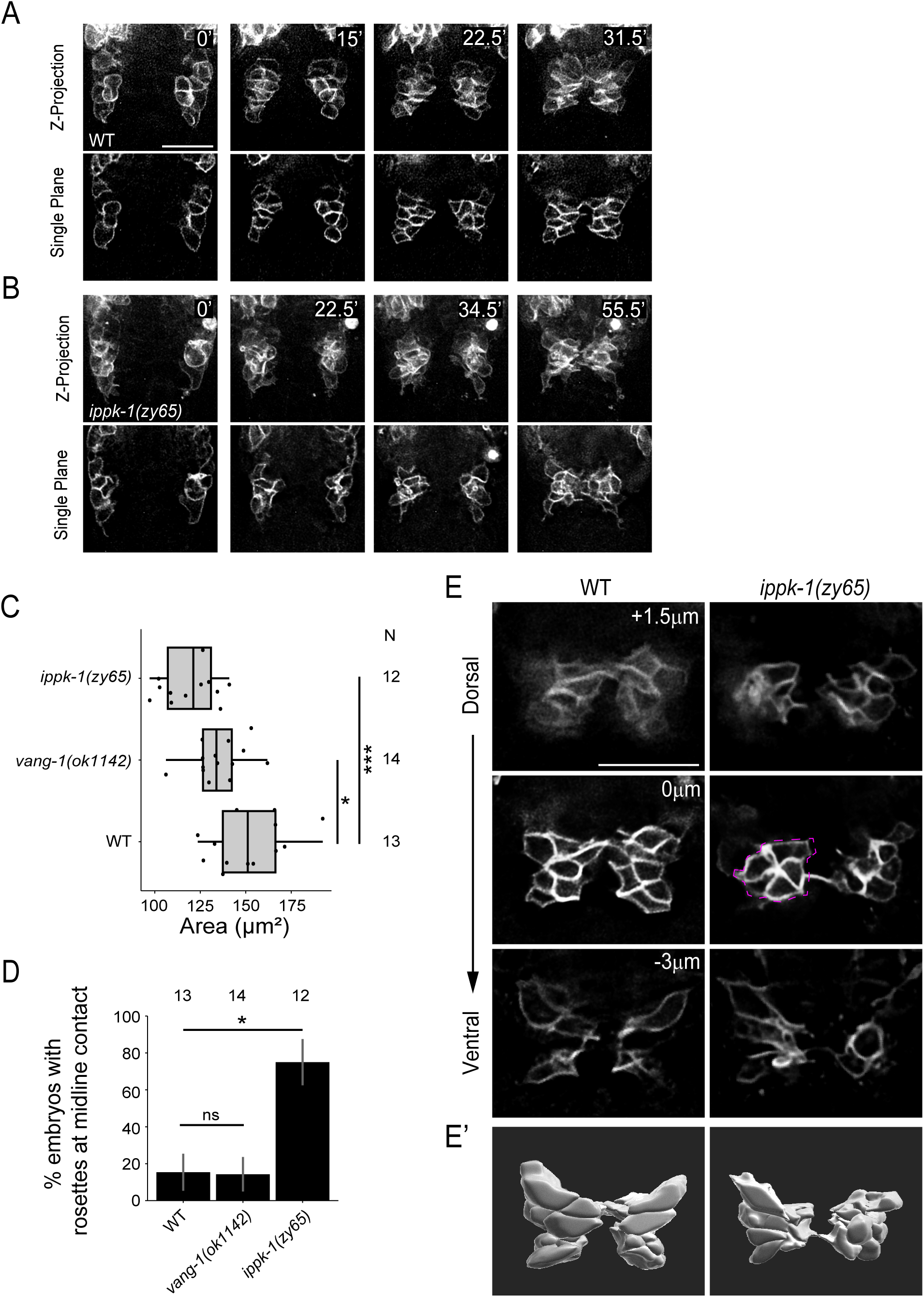
*ippk-1* mutants display defective organization of DD and DA neuronal progenitors prior to midline contact. (A-B) Time-lapse fluorescence single plane images and maximum-intensity projections showing the migration of a subset of DD and DA progenitors, labeled (membranes) with *cnd-1p::PH::mCherry*, from left and right sides toward the midline in (A) WT and (B) *ippk-1(zy65)* embryos. Scale bars = 10μm. (C) Box plot showing the combined area of left and right *cnd-1p*-labeled VNC progenitors at midline contact. Tails indicate min and max. (D) Plot showing the proportion of embryos containing at least one rosette among VNC progenitors at midline contact. Error bars indicate SEP. (E) Z-slices arranged from dorsal to ventral from a representative WT and *zy65* embryo showing *cnd-1p*-labeled VNC progenitors at midline contact, revealing the more compact organization and the presence of a rosette (dotted magenta outline) in the *zy65* embryo. (E’) 3D reconstruction of VNC progenitors in E. Statistics: (C) Chi-squared test with Monte Carlo simulation for 10,000 replicates, followed by a pairwise analysis using Fisher’s exact test with Monte Carlo simulation for 10,000 replicates and adjustments with Holm corrections, (D) ANOVA with Dunnett’s post-hoc test against WT. *p<0.05, **p<0.01, ***p<0.001.

To ascertain whether the more compact organization resulted from changes in cell number or fate, we counted the number of *cnd-1* promoter-labeled progenitors at the embryonic bean stage and in L1 animals using terminal cell fate markers. We found a slight reduction (p=0.0131) in the number of progenitors at the bean stage, with an average of 20 *cnd-1*-labeled cells (N=16) in WT and 19 (N=19) in *ippk-1* mutants (Fig. S5A). However, the number of terminally differentiated DD, DA, and DB neurons in L1 animals remained unchanged (Fig. S5B), suggesting that the more compact organization in *ippk-1* mutants is not due to a reduction in progenitor number or cell fate defects. Closer examination of the left and right progenitor groups to better understand their organization revealed differences in the presence of multicellular rosettes, structures where the junctions of five or more cells meet at a common vertex, between *ippk-1* mutants and WT. In *ippk-1(zy65)* mutants, 75% of embryos displayed a rosette at the time of midline contact (N=12, p=0.01294), compared to 15.4% in WT (N=13) embryos (Fig. 3D and E). These ectopic rosettes may therefore underlie some aspect of the disorganization and compactness seen in these progenitors during their migration to the midline. Ectopic rosettes appear to be a distinct feature of *ippk-1* mutants, as *vang-1(ok1142)* mutants did not show a greater proportion of rosettes at midline contact (N=14, p>0.9999) compared to WT (Fig. 3D). These observations suggests that IPPK-1 and its product IP6 are important for inhibiting rosette formation or promoting rosette resolution prior to VNC progenitor intercalation at the midline.

### *ippk-1* mutants display defects in convergent extension

Shortly after meeting at the midline, DD and DA progenitors from the left and right contribute to the formation of multicellular rosettes (Shah et al., 2017). The resolution of these rosettes sequentially from anterior to posterior is part of the convergent extension process, which narrows the tissue along the mediolateral axis and elongates it along the anterior-posterior axis. By the 1.5-fold stage, single-cell intercalations result in a stereotypical pattern of DD and DA neurons, aligned in a mostly single file along the developing VNC (Shah et al., 2017).

Our previous study indicated that the loss of PCP pathway components, like VANG-1, delays rosette resolution or causes unstable resolution, where rosettes partially resolve and then reform. These defects disrupt convergent extension and the normal intercalation of cells at the midline (Shah et al., 2017). To determine whether *ippk-1* regulates rosette dynamics, we performed time-lapse microscopy using our *cnd-1p::PH::mCherry* membrane marker to label DD and DA progenitors and measured the lifetime of the centrally positioned rosette formed when these cells meet at the midline. Rosette lifetimes were defined as the time from initial formation to resolution. For unstable resolutions, rosette lifetimes were measured from formation to final resolution. In WT, the centrally located rosette resolves within 3.3 minutes (median) after formation. In *ippk-1(zy65)* mutants, the median rosette lifetime increased to 17.1 minutes (p=0.0003) (Fig. 4A and B). As a comparison, we also measured rosette lifetime in *vang-1* mutants and showed that the median resolution time was 9.8 minutes (p=0.0082) (Fig. 4B). Interestingly, *vang-1(ok1142); ippk-1(zy65)* double mutants (17.2-minute median) did not show a significant increase in rosette lifetime compared to *zy65* single mutants (p>0.9999). This finding is consistent with *ippk-1* and *vang-1* acting in a common pathway to regulate rosette resolution dynamics.

**Figure 4.**
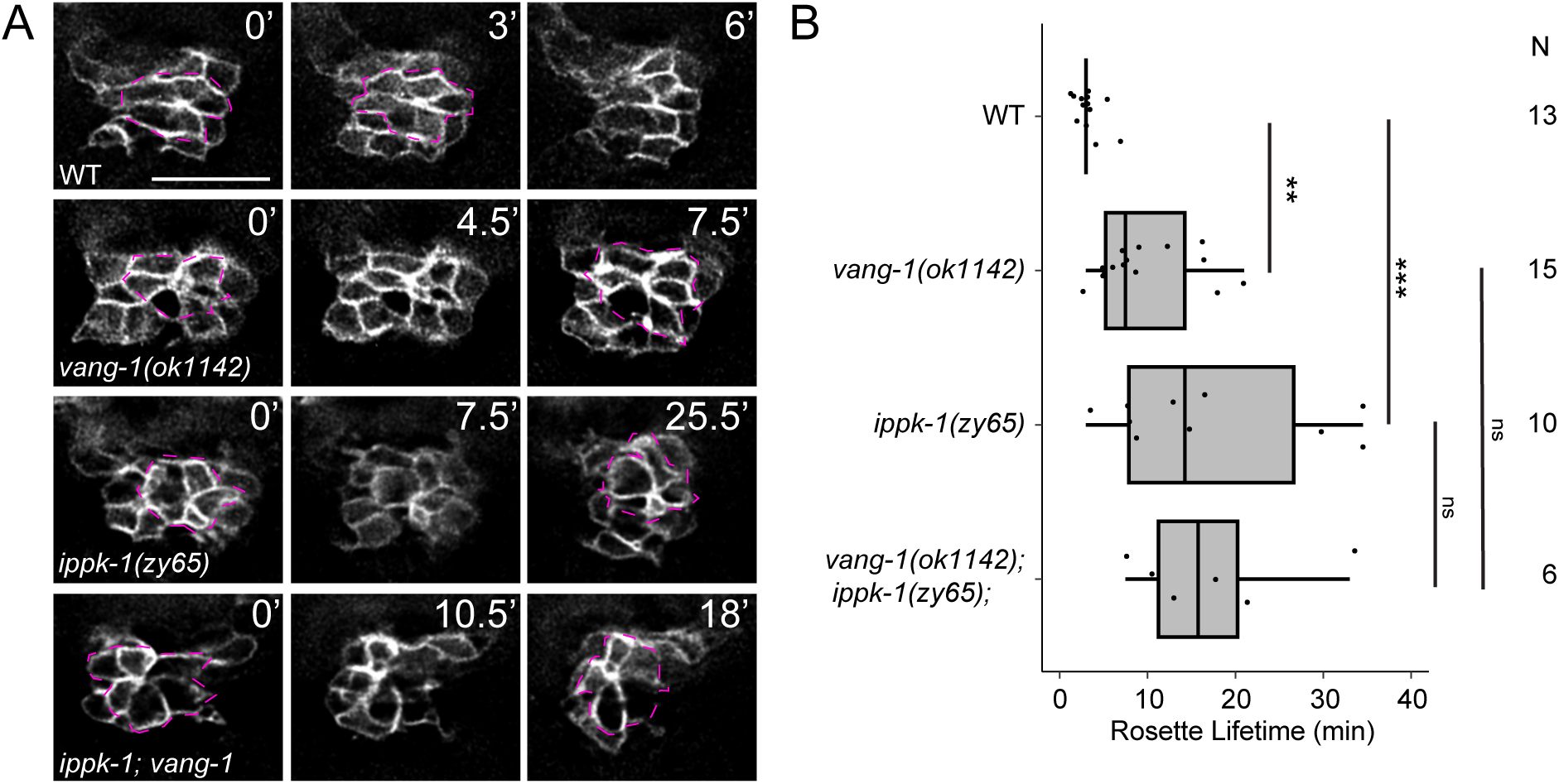
*ippk-1* mutants display defects in central rosette resolution. (A) Examples of rosette resolution dynamics in WT and mutant VNC progenitors labeled with *cnd-1p::PH::mCherry*. Cells participating in a rosette are outlined with a dotted magenta line. Mutant timelapses depict examples of unstable rosette resolution, where rosettes partially resolve and then reform. (B) Quantification of central rosette lifetime. Box plot tails indicate min and max. Statistics: Kruskal-Wallis with Dunn’s post-hoc test. *p<0.05, **p<0.01, ***p<0.001. Scale bars = 10μm.

The outcome of single cell intercalation and effects on VNC extension were assessed at the 1.5-fold stage using *unc-30p::GFP* to label DDs (cytoplasm) and *cnd-1p::PH::mCherry* to label DD and DAs (membrane). At this stage, most DD and DA progenitors have intercalated into a single file with some of DD1-4 and DA2-5 displaying a stereotypical alternating DD DA pattern (WT in Fig. 5A). We defined an embryo with a cell intercalation defect as one containing at least one abnormal DD-DD cell contact instead of a DD-DA contact. Not surprisingly, given the earlier organization and rosette resolution defects, we found a significant number of single cell intercalation defects in *ippk-1(zy65)* (36.5%) (N= 52, p=0.0069) and *ippk-1(or1572)* (50%) (N= 32, p=0.0004) mutants compared to WT (7.7%) (N=52) (Fig. 5A and B). *ipmk-1* mutants showed a modest increase at 28.2% (N=39, p=0.0912) in single cell intercalation defects compared to WT, although this difference was not statistically significant.

**Figure 5.**
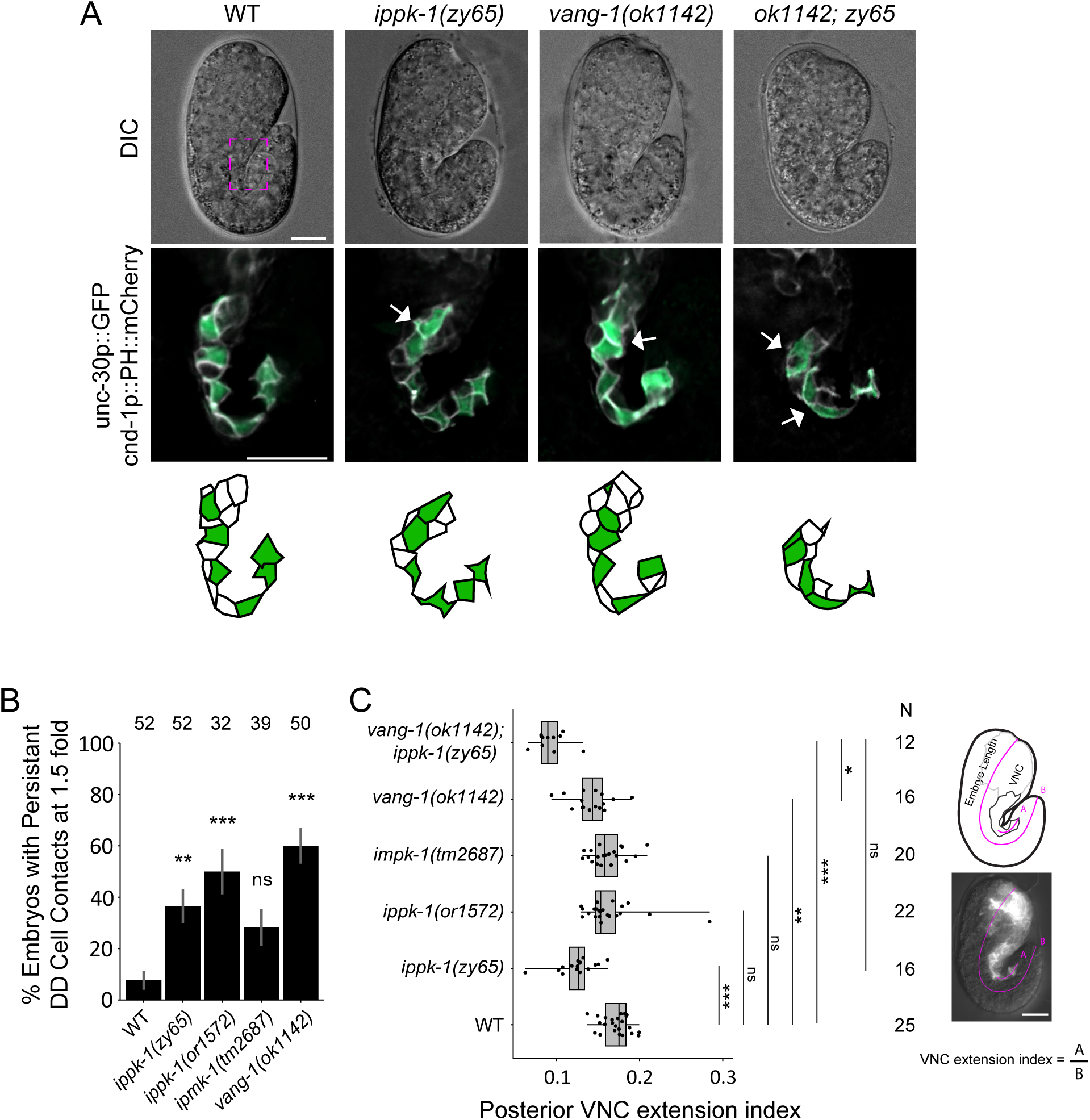
*ippk-1* mutants display neuron intercalation and VNC extension defects. (A) Brightfield images of embryos and corresponding deconvolved fluorescence images of the VNC at 1.5 fold in WT and mutants, with the cytoplasm of DD neurons labeled with *unc-30p::gfp* and the membranes of DD and DA neurons labeled with *cnd-1p::PH::mCherry*. The magenta box indicates the approximate area of the middle panels. Schematics of the corresponding middle panel VNC show DD neurons highlighted in green. (B) Quantification of persistent DD cell contacts in WT and mutant embryos. Error bars indicate SEP. Statistics: Chi-squared test, followed by a pairwise analysis using Fisher’s exact test adjusted with Bonferroni corrections. (C) Quantification of the posterior VNC extension index in WT and mutant embryos, relative to total embryo length. Tails indicate min and max. Panels show an image and corresponding schematic of a WT embryo with magenta lines indicating the lengths measured to calculate index. Statistics: (B) Chi-squared test with Monte Carlo simulation for 10,000 replicates, followed by a pairwise analysis using Fisher’s exact test with Monte Carlo simulation for 10,000 replicates and adjustments with Holm corrections, (C) Kruskal-Wallis with Dunn’s post-hoc test, significance compared to WT is shown. *p<0.05, **p<0.01, ***p<0.001. Scale bars = 10μm.

To determine if loss of *ippk-1* affected VNC extension at the 1.5-fold stage, we calculated a VNC extension index, defined as the length the VNC extends past the apex of the embryonic fold divided by the overall embryo length (Fig. 5C). In WT animals, this index is 0.17, whereas it showed a slight but significant decrease to 0.12 in *ippk-1(zy65)* mutants (p<0.0001), indicating a shorter posterior extension (Fig. 5C). However, neither *ippk-1(or1572)* nor *ipmk-1(tm2687)* mutants showed a significant change. Interestingly, within the scored *or1572* embryos, one exhibited an unusually long posterior extension, though the reason for this remains unclear. *vang-1* mutants also showed a small but significant decrease in the extension index to 0.14 (p=0.0069). In *vang-1; ippk-1* double mutants, this index decreased even further to 0.09 (p<0.0001) and was smaller than *vang-1* (p<0.0348) but not significantly different than *ippk-1* single mutants (p<0.9999) (Fig. 5C). This result is consistent with the rosette lifetime findings, suggesting that *ippk-1* and *vang-1* may function in a common pathway. Together, defects in rosette resolution, single cell intercalation and posterior VNC extension are consistent with the involvement of *ippk-1* in convergent extension during VNC assembly.

### Exogenous IP6 is sufficient to rescue *ippk-1* defects

In the IP kinase cascade, IPPK-1 phosphorylates IP5 to generate IP6. As a result, we explored whether the introduction of exogenous IP6 could mitigate the phenotypes associated with *ippk-1(zy65)*. To examine this, we performed microinjections of either IP6 (phytic acid) or a control solution (H_2_O) into the gonads of *ippk-1(zy65)* worms and subsequently evaluated their progeny for signs of rescued left and right embryonic DD DA organization and larval DD neuron cell body position defects. Measurement of IP6 levels in different cells and organisms suggested that endogenous concentrations ranged from 10µM to 100µM (Szwergold et al., 1987; Freund et al., 1992; Veiga et al., 2006). Consequently, we tested both 10µM to 100µM IP6 concentrations to assess their potential to rescue *ippk-1* mutants.

Injection of IP6 into *ippk-1(zy65)* mutant mothers restored the organization of *cnd-1*-labeled left and right-side DD and DA progenitors, as determined by scoring ectopic rosettes. *zy65* mutants with ectopic rosettes were reduced from 58.1% (N=31) to 11.8% (N=34) (p=0.0002) in F1 progeny of mothers injected with 100µM IP6 (Fig. 6A and B). Similarly, exogenous IP6 was able to rescue DD neuron positioning defects in L1 and L2 larvae. In WT, the six DD neurons, visualized with a *flp-13p::GFP* reporter, are typically equidistantly spaced along the VNC, whereas in *zy65* mutants, they are anteriorly displaced. For simplicity, we only assessed the spacing between DD1 and DD2, considering it defective if they were within one cell diameter of each other. In F1 progeny, the proportion of *zy65* mutants with DD1 DD2 spacing defects decreased significantly from 94.1% (N=51) to 37.9% (N=87) and 23.5% (N=68) when mothers were injected with 10µM or 100µM IP6, respectively (both p<0.0001) (Fig. 6C and D). Injection of IP6 into WT animals did not result in significant ectopic rosette or DD spacing defects in their progeny. As a specificity test, we injected 100µM IP6 into *vang-1* mutant mothers and observed no rescue of DD1 DD2 spacing defects in the F1 progeny (Fig. 6D). This indicates that exogenous IP6 specifically corrects defects associated with IPPK-1 function. These findings show that IP6, the downstream product of IPPK-1, is sufficient to rescue *ippk-1* defects, suggesting that these defects result from an IP6 deficiency.

**Figure 6.**
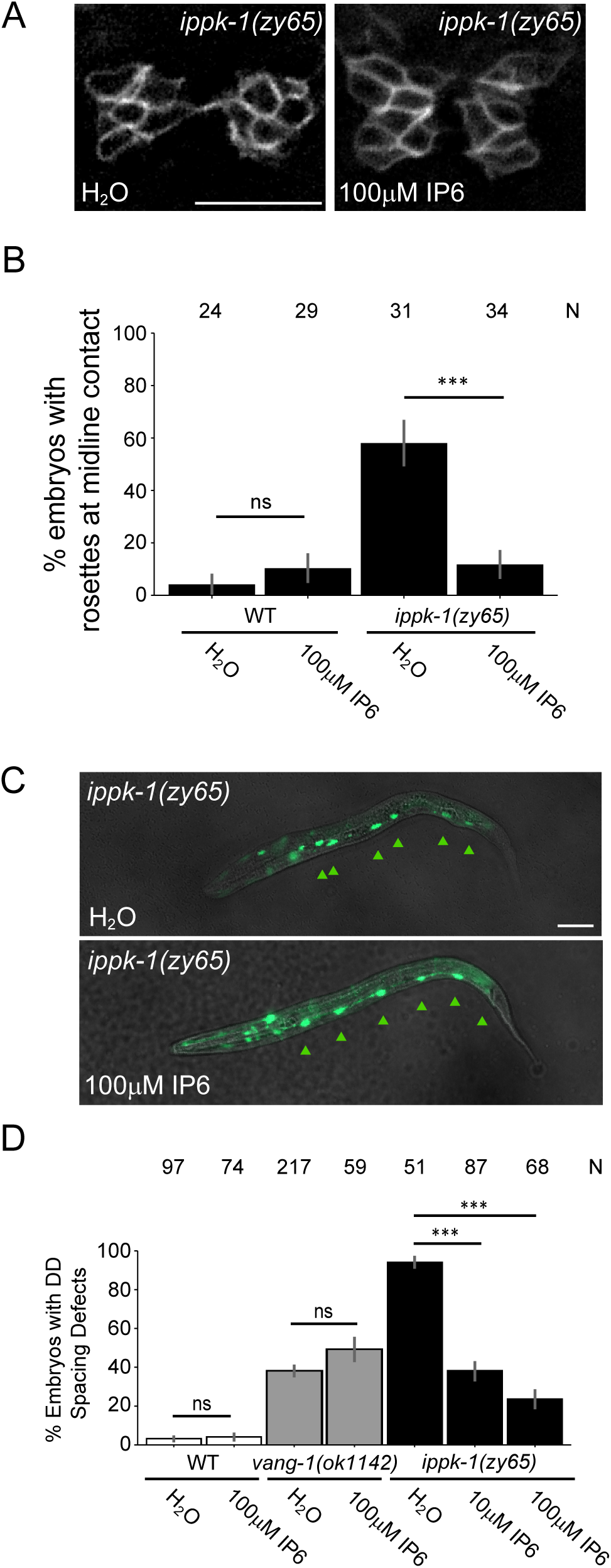
Exogenous IP6 is sufficient to rescue *ippk-1* VNC defects. (A) Representative images of VNC progenitors at the stage of midline contact in *ippk-1(zy65)* embryos, showing rescue of organization defects following injection with either H_2_0 or IP6 (dissolved in H_2_0). Scale bar = 10μm. (B) Plot showing the proportion of embryos in which VNC progenitors form at least one rosette at midline contact. (C) Representative images of DD neurons in *ippk-1(zy65)* larvae showing rescue of position defects after injection with IP6. Arrowheads mark DD neurons. Scale bar = 20μm. (D) Quantification of the proportion of WT and mutant larvae displaying a DD1 DD2 spacing defect. For both B and D, error bars indicate SEP. Statistics: (B) Fisher’s exact test. (D) Control and treatment groups for WT and *vang-1(ok1142)* were compared using Fisher’s exact test. *ippk-1(zy65)* strains were compared using a Chi-squared test with Monte Carlo simulation for 10,000 replicates, followed by a pairwise analysis using Fisher’s exact test with Monte Carlo simulation for 10,000 replicates and adjustments with Holm corrections.

## Discussion

IP molecules, through their regulation of basic cellular processes, play essential roles during animal embryonic development (Seeds et al., 2015; Frederick et al., 2005; Sarmah et al., 2005; Versky et al., 2005a). The pathway for IP synthesis involves a cascade of inositol kinases that sequentially phosphorylate one of six hydroxyl positions in the inositol ring. It typically commences with phospholipase C (PLC) and phosphoinositide phosphatidylinositol (4,5)-bisphosphate (PIP2)-derived IP3, ultimately leading to the production of IP6, as well as the higher inositol pyrophosphates IP7 and IP8 (Laha et al., 2021; Majerus, 1992). Inositol-pentakisphosphate 2-kinase (IPPK) is the rate limiting kinase that converts IP5 to IP6, by phosphorylating IP5 at the 2 hydroxyl position of the inositol ring (Verbsky et al., 2005b; Verbsky et al., 2002). In this study, we show that the *C. elegans* orthologue IPPK-1 and its product IP6 are important for proper ventral nerve cord (VNC) morphogenesis.

IPPK-1 in *C. elegans* has previously been shown to be essential for the formation and maintenance of germline membrane architecture (Lowry et al., 2015). We found that IPPK-1 is also important for multiple aspects of the morphogenetic process that leads to the narrowing and elongation of neuronal progenitor tissue as its constituent cells move towards and intercalate at the midline to form the VNC. The loss of *ippk-1* disrupts this process in several ways. First, the DD and DA progenitor assemblies arising on the left and right sides of the embryo are more compact and disorganized as they converge toward the midline. This abnormal tissue morphology may, in part, result from a significant increase in ectopic rosettes, structures where five or more cells meet at a common vertex, within these assemblies. Second, *ippk-1* mutants display delays in resolving the multicellular rosettes that form when the left and right VNC progenitors meet at the midline. Rosette dynamics are important drivers of convergent extension, as they reorient progenitors from the left and right to align along the anterior-posterior axis. And third, defects in midline intercalation that result in motor neuron cell body mispositioning along the developing VNC at the 1.5-fold stage of embryonic development. Collectively, these defects contribute to the anterior displacement of VNC motor neurons observed in *ippk-1* mutants at hatching.

In metazoans, IP kinases are classified into four groups: IPK, ITPK, IPPK, and PPIP5K, with the IPK group further subdivided into ITP3K, IPMK and IP6K (Laha et al., 2021). Some of these kinases, like ITPK and IPMK are multifunctional and phosphorylate more than one IP ring position (Odom et al., 2000; York et al., 1999). The *C. elegans* genome encodes one orthologue each of ITP3K, IPMK, IPPK and PPIP5K and three orthologues of IP6K. IPMKs catalyze the stepwise phosphorylation of the 6 and 3 hydroxyl positions of IP3 and IP4 respectively to generate IP5. Loss of the *C. elegans* IPMK (*ipmk-1)* leads to phenotypic traits like those caused by *ippk-1* loss, including intercalation defects and neuron position defects in newly hatched larvae. This finding, coupled with the rescue of *ippk-1* defects by the exogenous addition of IP6, indicates that VNC defects in *ippk-1* mutants are caused by disruption of IP metabolism and not some other factor independent of its catalytic activity.

Surprisingly, the defects in *ipmk-1* were generally milder than those of *ippk-1*, even though we used a deletion allele expected to result in a strong loss of function and, therefore, would be expected to strongly disrupt IP5 production. In vertebrates, the ability of ITPK, acting through an alternative PLC-independent pathway, to sequentially phosphorylate hydroxyl positions on IP1 through IP4 to generate IP5, could potentially compensate for the loss of IPMK (Desfougères et al., 2019). However, in contrast to vertebrates, *C. elegans* and several other lower species do not possess a member of the ITPK family (Laha et al., 2021). One possibility is that in *C. elegans*, another IP kinase has an IP binding pocket that enables it to bind and phosphorylate IP4 molecules. A possible candidate for such a multifunctional kinase might be LFE-2, the ITP3K orthologue. Supporting this idea, the simultaneous loss of both *ipmk-1* and *lfe-2* is lethal, even though each gene, when mutated individually, does not affect viability (Yang et al., 2021).

In a previous study, we found that VANG-1 and PRKL-1, core components of the planar cell polarity pathway (PCP), and SAX-3, a Robo receptor, act in parallel pathways to regulate convergent extension during VNC assembly (Shah et al., 2017). Disruption of these pathways individually leads to cell intercalation defects and prolonged rosette resolution times, which may contribute to the anterior neuron displacement observed in the mutants. However, simultaneous loss of both pathways results in a more severe disruption of rosette-mediated convergent extension. For example, in our previous study, we found that in wild type, the central rosette that forms at the midline resolves in less than 4 minutes, compared to approximately 20 minutes in *sax-3* and *prkl-1* single mutants and greater than 50 minutes in *sax-3; prkl-1* double mutants (Shah et al., 2017). This long delay in the double mutant before midline intercalation, during which time the embryo continues to undergo elongation into a more wormlike shape, likely explains why, at hatching, nearly all motor neuron cell bodies are abnormally positioned in the anterior half, rather than being evenly distributed along the VNC.

Loss of *ippk-1* results in defects similar to those of *vang-1* mutants, including rosette resolution delays, incorrect cell contacts following midline intercalation, and anterior displacement of motor neurons in newly hatched worms. We found that the average rosette delay phenotype of *vang-1; ippk-1* double mutants was not significantly different from that observed in the single mutants. This suggests that *ippk-1* and *vang-1* may act within a common pathway to ensure timely rosette resolution. However, the involvement of independent pathways cannot be excluded, as the *ippk-1* mutants used in the double mutants likely retain substantial IP5 kinase activity, given that complete loss of function is lethal. Consequently, the full extent of *ippk-1* involvement in VNC assembly remains unclear and we cannot rule out the possibility that a complete lack of IPPK-1 leads to the severe disruption of convergent extension observed in PCP and Robo double mutants.

Interestingly, despite similar rosette lifetimes in embryos, *vang-1; ippk-1* double mutants exhibited more pronounced anterior shifts in DD neuron cell body positions in larvae compared to single mutants. This observation suggests that proper neuronal positioning in the VNC may rely on both rosette-dependent and rosette-independent mechanisms. Indeed, much remains to be understood about VNC assembly, particularly the molecular and cellular processes that ensure the proper spacing and stereotypical organization of the three motor neuron classes, DD, DA, and DB, present in the VNC of L1 larvae.

The role that IPPK-1 plays in VNC development is interesting given the evidence highlighting the importance of IP metabolism in neural tube formation. Several studies have shown that disruptions in IP kinases are associated with neural tube defects. Notably, loss of mammalian ITPK1 leads to embryos that develop neural tube defects of varying severity, including spina bifida and exencephaly (Wilson et al., 2009). A similar pattern is observed in mutants for the mammalian IPMK/IPK2; although embryos arrest around E9.5, they exhibit multiple morphological abnormalities, including abnormal folding of the neural tube (Frederick et al., 2005). ITPK polymorphisms have been associated with an increased prevalence of neural tube defects and are correlated with lower maternal plasma IP6 levels (Guan et al. 2014). Similarly, reduced maternal myo-inositol levels are associated with an increased risk of spina bifida (Groenen et al., 2003). However, IPPK has not been directly linked to neural tube defects, as loss of *Ippk* in mouse models leads to early embryonic defects that prevent a thorough analysis of neural tube development (Verbsky et al., 2005a). It is important to note, though, that all three genes are expressed in the developing neural tube (Wilson et al., 2009; Frederick et al., 2005; Versky et al., 2005a), consistent with their shared role in IP metabolism.

Convergent extension is a key driver of neural tube formation in chordate species. In mice, mutations in genes associated with PCP, which are key regulators of convergent extension, are closely linked to failed neurulation (Lesko et al., 2021; Williams et al., 2014; Blankenship et al., 2006). Convergent extension and neural tube development in mice are, in part, driven by the regulation of junctional dynamics, including the formation and resolution of multicellular rosettes (Williams et al., 2014). In other animals, convergent extension has been shown to depend on localised junctional dynamics, which require the asymmetric distribution of PCP proteins (Shindo et al., 2019; Butler and Wallingford, 2018). IP6 and its derivatives have been previously shown to interact with several membrane and cytoskeletal proteins both *in vitro* and *in vivo* during cell migration and adhesion (Cheng and Andrew, 2015; Fu et al., 2017; Schröterová et al., 2018; Rojas et al., 2019; Bhat et al., 2024). Given the importance of cytoskeleton dynamics in collective cell movements, the production of these metabolites may play a role in facilitating membrane dynamics. Supporting this idea, loss of *ippk-1* disrupts the complex membrane architecture of the germline, which relies on membrane remodeling during oogenesis (Lowry et al., 2015).

Another potential link between IP6 and convergent extension is the septin family, a group of cytoskeletal proteins involved in membrane dynamics and cytoskeletal organization. Septins have been implicated in the establishment of polarized actin networks during collective cell behaviors, including during convergent extension in *Xenopus* mesoderm (Devitt et al., 2024). Septin proteins co-localize and function alongside PCP components, including Vangl2 and Prickle2, influencing the distribution of actin rich cables along the anterior-posterior axis. An IP6 affinity tag is also able to bind with septin proteins in a colon cancer cell line alongside intercellular transport and cytoskeleton associated targets (Yin et al., 2016).

The production of IP6 may also facilitate convergent extension more broadly through the regulation of endocytosis. Several types of inositol phosphates and pyrophosphates bind to components of vesicle transport pathways, including AP-1/2/3, PIP2, synatotagmin-1, and dynamin, where they facilitate both exocytosis and endocytosis in diverse situations (Li et al., 2022; Yang et al., 2012; Høy et al., 2002; Voglmaier et al., 1992). Endocytosis is essential for convergent extension movements, as evidenced by defects in intercalation and tissue extension when dynamin-mediated endocytosis is disrupted (Levayer et al., 2011; Jarrett et al., 2002). While endocytosis is a common feature of numerous signalling pathways, one context in which it plays a significant role is in the regulation and establishment of PCP signalling through mechanisms involving protein trafficking (Ohkawara et al., 2011; Onishi et al., 2013; He et al., 2018). Multiple regulators of convergent extension during gastrulation for example, including the secreted Frizzled-related protein 2 (sFRP2) and phospholipase D (PLD1) in *Xenopus* and the E3 ubiquitin ligase Mindbomb1 in Zebrafish, either promote or redirect PCP signalling through internalization of Frizzled or the co-receptor Ryk (Brinkmann et al., 2016; Lee et al., 2016; Saraswathy et al., 2022). Similarly, the PCP pathway may interact with other signaling pathways via endocytosis, as demonstrated by Disheveled 2-mediated endocytosis of the EphrinB receptor during Zebrafish neurulation (Kida et al., 2007).

Overall, our findings implicate IPPK-1 and its metabolite IP6 in coordinating the organization and timing of cell intercalations during VNC assembly. Together with the involvement of IP kinases in vertebrate neural tube formation, this suggests that IP metabolism plays an evolutionarily conserved role in central nerve cord development. Considering the crucial role of convergent extension in this process, it will be important to elucidate the precise mechanism through which IP6 or other inositol phosphates regulate this morphogenetic event.

## Lead contact

For further information and requests for resources and reagents, contact Antonio Colavita.

## Materials and methods

### Key resources table

| Reagent or Resource | Source | Identifier |
| --- | --- | --- |
| Chemicals, Peptides, and Recombinant Proteins |  |  |
| Phytic acid sodium salt hydrate | Sigma-Aldrich | P0109; CAS:14306-25-3 |
| Levamisol hydrochloride | Millipore Sigma | 31742; CAS:16595-80-5 |
| **Bacterial Strains** |  |  |
| *E. coli*: OP50 | *Caenorhabditis*<br>Genetics Center<br>(CGC) | WBStrain00041969 |
| **Experimental Models: Organisms/Strains** |  |  |
| *C. elegans*: N2 (wild isolate) | CGC |  |
| *C. elegans*: *ippk-1*(zy65) II | This study |  |
| *C. elegans*: *ippk-1*(or1572) II | CGC, Lowry et al.,<br>2015. | EU2908<br>WBVar00241916 |
| *C. elegans*: *ippk-1*(zy100[*GFP::SEC::IPPK-1*]) II | This study |  |
| *C. elegans*: *ippk-1*(zy102[*IPPK-1::GFP*]) II | This study |  |
| *C. elegans*: *ynIs37*[*flp-13p::gfp*] III | CGC | NY2037<br>WBStrain00029158 |
| *C. elegans*: *ipmk-1*(tm2687) IV | Shohei Mitani,<br>National<br>BioResource Project<br>(NBRP) |  |
| *C. elegans*: *sax-3*(zy5) X | Shah et al., 2017 |  |
| *C. elegans*: *vang-1*(ok1142) X | CGC | RB1125<br>WBStrain00031829 |
| *C. elegans*: *zyIs36*(*cnd-1::PH::mCherry*; *myo-2p::mCherry*) X | Antonio Colavita,<br>Shah et al., 2017 | RW12267 |
| *C. elegans*: *zyIs40*(*unc-30p::gfp*; *cnd-1::PH::mCherry*) Unassigned linkage | This study |  |
| *C. elegans*: *vab-7*(*st12267*[*vab-7::GFP*]) | CGC |  |
| **Plasmids** |  |  |
| pPD95.77 | Andrew Fire | Addgene<br>#1000000001 |
| pDD282 | Dickinson et al.,<br>2015 | Addgene #66823 |
| pDD162 | Dickinson et al.,<br>2013 | Addgene #47549 |
| pAC422 | This study |  |
| pAC425 | This study |  |
| pAC426 | This study |  |
| pCFJ90 | Frøkjær-Jensen et<br>al., 2008 | Addgene #19327 |
| **Oligonucleotides** |  |  |
| DNA Oligonucleotides | Eurofins | See Table S1 for<br>sequences. |
| **Software and Algorithms** |  |  |
| Zeiss Zen 3.2 | Zeiss | https://www.zeiss.com/microscopy/en/products/software/zeiss-zen.html |
| Graphpad Prism 10 | Graphpad Software | https://www.graphpad.com/scientific-software/prism/ |
| Imaris 10 | Oxford Instruments | https://imaris.oxinst.com/ |

### Strains and culture conditions

Strains were maintained on NGM plates at 20°C, except for temperature-sensitive strains, which were maintained at 18⁰C. The Bristol N2 strain was used as wild type (WT), along with the following alleles and transgenes: LGII: *ippk-1(zy65), ippk-1(zy100[GFP::SEC::ippk-1]), ippk-1(zy101[GFP::ippk-1]), ippk-1(zy102[ippk-1::GFP]), ippk-1(or1572), unc-4(syb1658[unc-4::GFP]).* LGIII: *ynIs37[flp-13p::gfp], vab-7(st12267[vab-7::GFP])*. LGIV: *ipmk-1(tm2687).* LGX: *sax-3(zy5), vang-1(ok1142), zyIs36[cnd-1p::PH::mCherry myo-2p::mCherry].* Unassigned linkage: *zyIs40[unc-30p::GFP; cnd-1p::PH::mCherry myo-2p::mCherry]*.

### Whole genome sequencing and GFP insertion into the endogenous *ippk-1* gene

*zy65* was identified in a genetic screen for DD neuron position defects (A Colavita, unpublished) and outcrossed at least 3 times. The mutation in the *zy65*-containing strain was identified by whole genome sequencing as described in Noblett et al. (2019). Sanger sequencing (performed at OHRI Stemcore) was used to genotype *zy65* in single and compound mutants, using the primers N22 and N23 (Table S1).

Two separate strains were generated in which GFP was inserted at either the N-terminus or the C-terminus of the endogenous *ippk-1* gene using the CRISPR/Cas9 homology-directed repair (HDR) approach described in Dickinson et al. (2015). Target sites for N-terminal and C-terminal insertion were generated using the IDT guide RNA design tool at idtdna.com/site/order/designtool/index/CRISPR_SEQUENCE. Guide sequences were inserted into the Cas9–sgRNA plasmid (pDD162, Addgene #47549) using the NEB Q5 Site-Directed Mutagenesis Kit. The HDR plasmid containing *ippk-1* homology arms and the GFP and selection cassette (pDD282, Addgene #66823) was made using Gibson Assembly as described in Dickinson et al. (2015). All primers, including sgRNA primers, are listed in Table S1.

### Reporter and rescue constructs

*ippk-1p::GFP* (pAC420) was made by amplifying a 2974 bp fragment of the *ippk-1* promoter from N2 genomic DNA using primers 18 and 19 (Table S1) and inserting it between the *SphI* and *BamHI* sites of pPD95.77 (from Andrew Fire Lab *C. elegans* Vector Kit). GFP::IPPK-1 was made using Gibson Assembly (New England Biolabs) by combining fragments from cDNA clones yk1067e07 and yk1209d01 (kindly provided by Yuji Kohara, National Institute of Genetics, Japan) and pPD95.77 using primers N12-N17 (Table S1). The resulting construct (pAC422) contained a 2246 bp *ippk-1* cDNA, which was comprised of a 1365 bp open reading frame and an 881 bp 3’ untranslated region, inserted in-frame with GFP. The ippk-1p::GFP::IPPK-1 construct (pAC426) was made by amplifying the 2974 bp *ippk-1* promoter with primers N18 and N19 (Table S1) and inserting it between the *SphI* and *BamHI* sites of pAC422. The unc-33p::GFP::IPPK-1 construct (pAC425) was made by amplifying the 1961 bp *unc-33* promoter from N2 genomic DNA using primers N20 and N21 (Table S1) and inserting it between the *SphI* and *BamHI* sites of pAC422. Germline transformation was performed using standard methods (Mello et al., 1991) by injecting constructs at 1-5 ng/µl along with 2.5 ng/µl of *myo-2p::mCherry* (pCFJ90, Addgene #19327) and pBluescript KS (Stratagene) to a total DNA concentration of 100 ng/µl.

### Microscopy and image analysis

For static and timelapse imaging of live embryos, L4 worms were grown at 25°C, except for *ippk-1(or1572)* which was grown at 18°C, until gravid. Static imaging was used to quantify the posterior extension of the VNC, presence of persistence-cell contacts and to determine the number of *cnd-1* positive VNC progenitors. Embryos were imaged at 63x magnification using a Zeiss Imager M2, Colibri7 LED source, AxiocamHR camera and Zen V3.2 software.

Tissue measurements were taken with a z-projection of 42 slices with 0.25μm slice width, in a *zyIs36 [cnd-1p::PH::mCherry]* background. Images were processed using a maximum intensity projection prior to analysis. The length of each embryo was measured down the midline starting at the tip of the mouth to the end of the tail. The posterior portion of the VNC was measured starting at the point where it transected the intestinal valve to the end of DD6 (Martin et al., 2016). All images were acquired with an RFP exposure time of 700-900ms at 30% intensity. Images for cell determination counts were taken with the same conditions across the width of the VNC using a 0.5μm slice width. All *cnd-1* positive neurons within the area of the VNC were counted. A representative image of the midline defects was also taken using these settings (Fig. 4C and D) and deconvolved using the constrained iterative algorithm on Zeiss Zen V3.2. Persistent-cell contact images were taken in a *zyIs40 (unc-30p::GFP cnd-1p::PH::mCherry)* background, with a GFP exposure time of 50ms at 15% intensity and mCherry exposure as above. Images were taken over a z-projection of 1-15 slices with 0.5μm slice width. A persistent cell contact was defined as the failure of WT intercalation between DD1-DD4, in at least two neurons.

Embryos used for time-lapse imaging were collected from plates using a capillary attached to glass pipette, mounted in a 3 μL drop of M9 containing approximately fifty 25 μm diameter polystyrene beads (Polyscience Inc.) and sealed under a coverslip using Vaseline. Fluorescence time-lapse imaging was performed at 63x magnification using the above microscope configuration with a Zeiss SVB1 microscope signal distribution box/*TTL* trigger cable-controlled camera shutter to minimize excitation light induced phototoxicity. Images of bean stage embryos were acquired at 1.5-minute intervals, for approximately 60-minutes, across a z-projection of 5 to 15 slices with 0.5 μm between slices. 2×2 binning was used to further minimize the required exposure times during timelapse for rosette timing measurements. Timelapse images were taken in a *zyIs36 [cnd-1p::PH::mCherry]* background, with an *mCherry* exposure time of 400ms at 10% intensity. The images were deconvolved using the iterative constrained algorithm on Zen 3.2. Measurements of VNC area were preformed on maximum intensity projections of the deconvolved images. Rosette duration was determined as performed in Shah et al. (2017). Images included in each figure were processed using Zeiss Zen V3.2 and Photoshop. Adjustments were made to the brightness-contrast of each image to standardize representative phenotypes to be easily visible. Deconvolution, using the iterative constrained algorithm on Zen 3.2, was preformed on a representative image of WT and *ippk-1(zy65)* VNC cells at the midline. 3D reconstructions were made by manually tracing cell outlines in Imaris (Oxford Instruments).

### Quantification of neuron number and position in larvae

Eight to ten L4 stage larval worms were transferred to seeded NGM plates and grown at 25°C overnight with the exception of *ippk-1(or1572)*, which was grown at 18°C. L1 stage progeny from these plates were mounted on 2% agarose pads using 200 μM levamisole (31742, Sigma). DD and DB neuron positions were measured as described in Shah et al. (2017). DD neurons were visualized with *ynIs37[flp-13p::GFP]* and DB neurons with *vab-7(zy142[vab-7::mNG::T2A::mScarlet-I::H2B])* (Saharkhiz et al., 2024). Briefly, neurons were scored by measuring the position of each along the worm using the curve (spline) tool (Zen 3.2). These positions were then converted into mean percentage locations using or DD1 or DB1 as 0% and the anus as 100%. Each percentage location was plotted using base R, version 4.4.2 (R Core Team, 2019) and geom_jitter from the ggplot2 suite (Wilkinson, 2011).

Individual DD, DA and DB neuron counts were scored at 20x magnification using *ynIs37[flp-13p::gfp]* (a gift from Dr. Chris Li, CUNY), *unc-4(syb1658[unc-4::GFP])* and *vab-7(st12267[vab-7::EGFP]* reporters respectively. Zeiss Zen 3.2 was used for image analysis.

### Exogenous IP6 rescue

IP6 (phytic acid sodium salt hydrate, P8810, Sigma) was dissolved in ddH_2_0 to prepare 0.1μM, 10μM and 100μM solutions. These concentrations were chosen, as they cover a range which has been quantified from mammalian and non-mammalian sources (Szwergold et al., 1987; Freund et al., 1992; Veiga et al., 2006). WT, *ippk-1(zy65)* and *vang-1(ok1142)* worms were grown at 18°C (*zy65*) or 20°C (WT and *ok1142*) and transferred to 25°C at 5-6 hours prior to injections. Young adult worms were then injected with IP6 solution into both gonad arms and allowed to recover for 1 hour. Healthy survivors were singled onto seeded NGM plates overnight at 25°C. Hatched larvae or embryos were mounted on the subsequent morning on 2% agarose pads. L1 and L2 stage worms were scored for the proximity of two or more neurons within a distance of one cell body. Ventral mounted embryos were scored for a shift in the initial meeting of cells towards the posterior of the embryo (as described above). Progeny from independent worms were pooled for statistical analysis.

### Statistics

Larval and embryo measurements were compared across populations with the indicated number of individuals. Statistical tests and parameters used to determine significance are indicated in each figure legend. All non-categorical data was assessed for normality when determining the appropriate statistical tests. If the datasets being tested passed the Shapiro–Wilk test, they were analyzed using either Welch’s T-test or analysis of variance (ANOVA), as appropriate. Groups that failed to pass the test were compared using the Mann-Whitney U test or Kruskal-Wallis test, as appropriate. Graphpad Prism 10 was used for statistical analysis, with the exception of Chi-squared tests followed by post hoc analysis which used base R, version 4.4.2 (R Core Team, 2019).

## Acknowledgments

We thank Shohei Mitani for sharing *ipmk-1(tm2687)* and Chris Li for sharing *ynIs37*. We also thank Saber Saharkhiz for comments on the manuscript. Some strains were provided by the *Caenorhabditis* Genetics Center, which is funded National Institutes of Health Office of Research Infrastructure Programs (P40 OD010440). Gene information, including genomic data, was accessed through WormBase. The authors acknowledge the Cell Biology and Image Acquisition Core (RRID: SCR_021845) funded by the University of Ottawa, Ottawa, Natural Sciences and Engineering Research Council of Canada, and the Canada Foundation for Innovation. We would also like to acknowledge the assistance of StemCore Laboratories Genomics Core Facility (OHRI, uOttawa), RRID:SCR_012601. This work was supported by grants from the Canadian Institutes of Health Research (123513 and 156160) to A. Colavita.

## Appendix A: Supplemental data

**Supplemental Figure 1.** i***p***pk***-1* mutants show defects in DB neuron positioning.** (A) Representative images of DB neuron positions in WT and *ippk-1* mutants, visualised using the DB-specific *vab-7(zy142[vab-7::mNG::T2A::mScarlet-I::H2B])*. Arrowheads mark DB neurons. Scale bar = 20μm. (B) Quantification of the mean position of DB1-DB7 neurons relative to DB1 in L1 stage WT and mutant worms. Neurons (colour coded as indicated along top) are plotted along the AP axis, where DB1 and anus mark the 0% and 100% positions respectively. Means and 95% confidence intervals are indicated for each DB2-7 neuron. Animals scored (N) are indicated on the right. Statistics: Two-tailed T-test was used, Welch’s correction. *p<0.05, **p<0.01, ***p<0.001. Arrows show the shift from WT for all neurons where p>0.05.

**Supplemental Figure 2.** R**e**scue **of *ippk-1(zy65)* DD neuron position defects with *ippk-1* and *unc-33* promoter driven *ippk-1*.** (A) Schematic of the transgene (TG) rescue experiment showing parental (P) and first (F1) generations, along with the presence or absence of TG maternal rescue (M^TG+^ or M^TG-^). (B) Representative images of DD positions in *ippk-1(zy65)* mutants with or without an extrachromosomal array carrying the IPPK-1 cDNA transgene and a *myo-2p::mCherry* pharyngeal muscle marker. DD neurons (arrowheads) are labeled using the DD-specific *flp-13p::GFP* reporter. Scale bar = 20μm. (C-D) *ippk-1(zy65)* DD position defects are maternally rescued when *ippk-1* is expressed from an *ippk-1* or pan-neuronal *unc-33* promoter. Quantification of DD2–4 positions relative to DD1 in L1 worms of the indicated genotype. Neurons (colour coded as indicated along top) are plotted along the AP axis, where DD1 and anus mark the 0% and 100% positions respectively. Means and 95% confidence intervals are indicated for each DD2-6 neuron. Animals scored (N) are indicated on the right. Statistics: One-way ANOVA with Dunnett’s post-hoc test against the corresponding WT neuron. *p<0.05, **p<0.01, ***p<0.001. Arrows show the size of the shift from the mean DD neuron positions found in a TG-*ippk-1(zy65)* mutant for all neurons where p>0.05.

**Supplemental Figure 3.** E**p**idermal **morphology defects in *ippk-1(zy65)* mutants.** (A-B) Representative image of (A) WT and (B) an *ippk-1(zy65)* embryos with epidermal protrusions (arrows). Scale bar = 10µm. (C-D) Representative image of (C) WT and (D) an *ippk-1(zy65)* larvae with epidermal protrusions (arrows). Scale bar = 20µm. (E) Quantification of epidermal defects in WT and mutant embryo and larvae.

**Supplemental Figure 4.** i***p***pk***-*1 expression.** (A) Schematic showing the *ippk-1* GFP transcriptional reporter and GFP knock-ins into the endogenous *ippk-1* gene. (B-D) Representative images of expression from an *ippk-1p::GFP* transgene containing ∼3kb of promoter sequence. Expression is detected in (B) pharynx, (C) spermatheca, and (D) tail cells. Endogenous *ippk-1* expression in adults from (E) *zy100[GFP::SEC::IPPK-1]* and (F) *zy102[IPPK-1::GFP]* showing strong expression in spermatheca (white arrows). Corresponding Nomarski image above fluorescence image. Scale bar = 20μm.

**Supplemental Figure 5.** i***p***pk***-1* mutants do not affect VNC progenitor numbers or terminal cell fates.** (A) Quantification of the number *cnd-1p::PH::mCherry* expressing VNC progenitors in late bean stage WT and *ippk-1(zy65)* mutants. (B) Quantification of the number of DA, DB and DD neurons in L1 WT and *ippk-1(zy65)* mutants. Statistics: (A) Mann-Whitney U test. (B) The number of larval DD and DA neurons were identical between WT and *ippk-1(zy65)* and thus no p-value can be provided. *p<0.05.

**Supplemental Table 1: Oligonucleotides used in this study.**

## Data availability

Data will be made available upon request.

