## Supplemental Figures S1-S5 for "IPPK-1 and IP6 contribute to ventral nerve cord assembly in *C. elegans*"

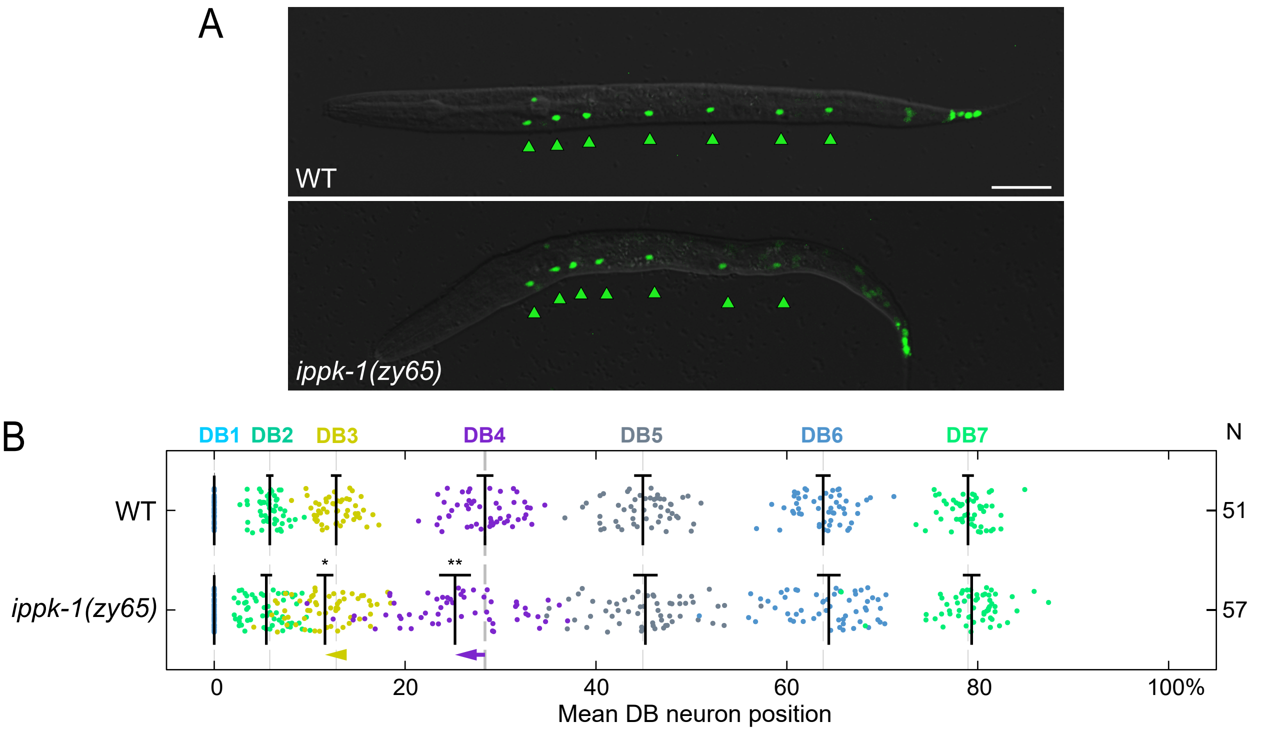
A
WT
ippk-1(zy65)
B
DB1 DB2 DB3 DB4 DB5 DB6 DB7
N
WT
51
ippk-1(zy65)
57
*
**
0
20
40
60
80
100%
Mean DB neuron position

**FIG S1**

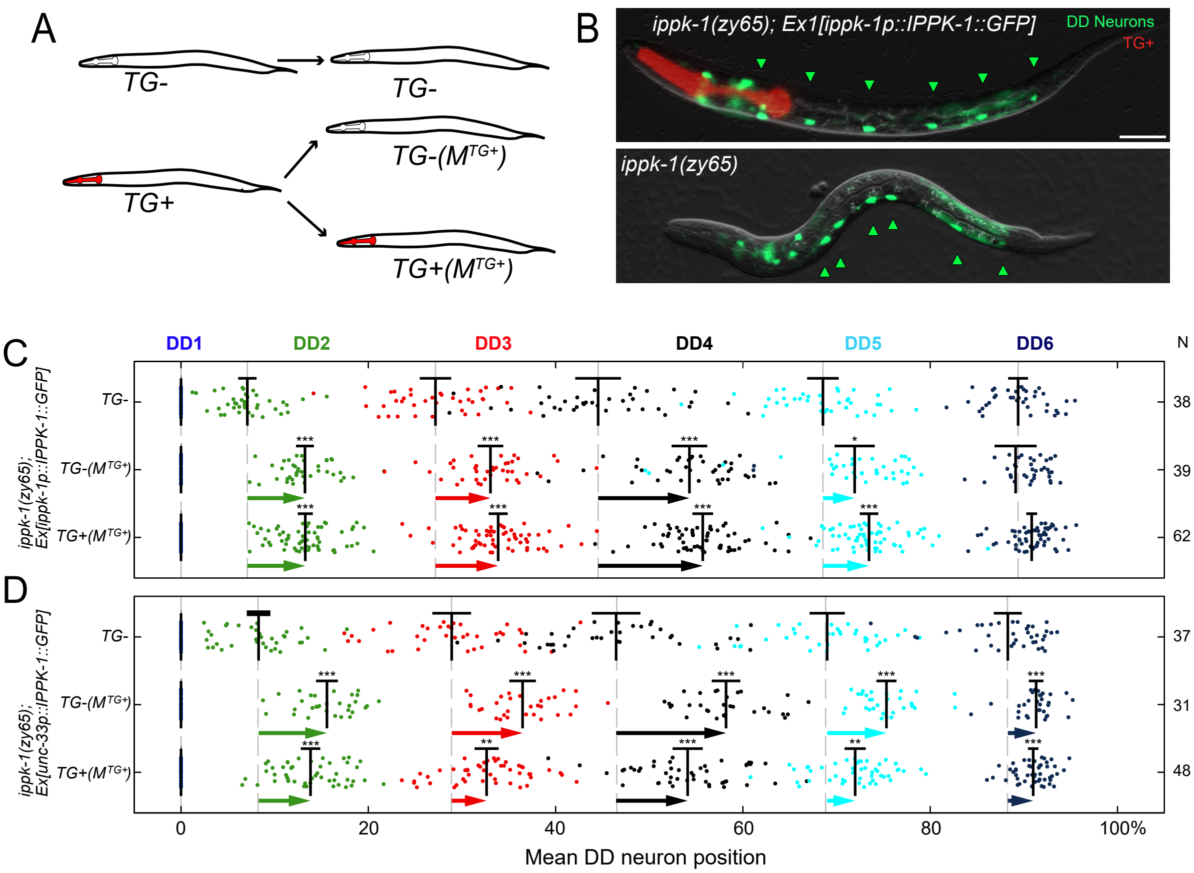
A
TG-
TG-
TG-($M^{TG+}$)
TG+
TG+($M^{TG+}$)
B
ippk-1(zy65); Ex1[ippk-1p::IPPK-1::GFP]
DD Neurons
TG+
ippk-1(zy65)
C
DD1
DD2
DD3
DD4
DD5
DD6
N
ippk-1(zy65); Ex[ippk-1p::IPPK-1::GFP]
TG-
TG-($M^{TG+}$)
TG+($M^{TG+}$)
38
39
62
D
ippk-1(zy65); Ex[unc-33p::IPPK-1::GFP]
TG-
TG-($M^{TG+}$)
TG+($M^{TG+}$)
37
31
48
0
20
40
60
80
100%
Mean DD neuron position

FIG S2

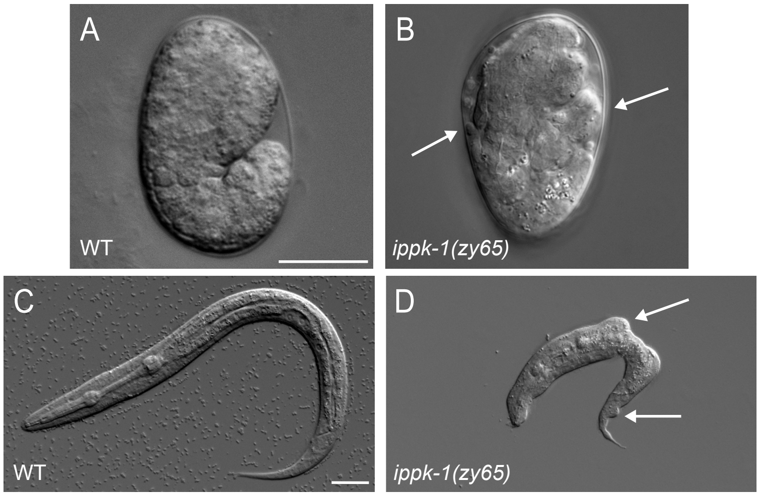


E

| **Mutant** | **# embryos (N)** | **% embryos with epidermal protrusions** | **# larvae (N)** | **% larvae with epidermal protrusions** |
|---|---|---|---|---|
| WT* | 76 | 0 | 79 | 0 |
| *ippk-1(or1572)** | 79 | 38.0 | 89 | 32.6 |
| WT | 75 | 0 | 65 | 0 |
| *ippk-1(zy65)* | 116 | 9.5 | 135 | 17.4 |
| *vang-1(ok1142)* | 93 | 0 | 67 | 0 |
| *ippk-1(zy65);vang-1(ok1142)* | 128 | 23.4 | 136 | 38.2 |
| *sax-3(zy5)* | 99 | 5.0 | 97 | 9.3 |
| *ippk-1(zy65);sax-3(zy5)* | 113 | 32.7 | 153 | 60.8 |

**FIG S3**

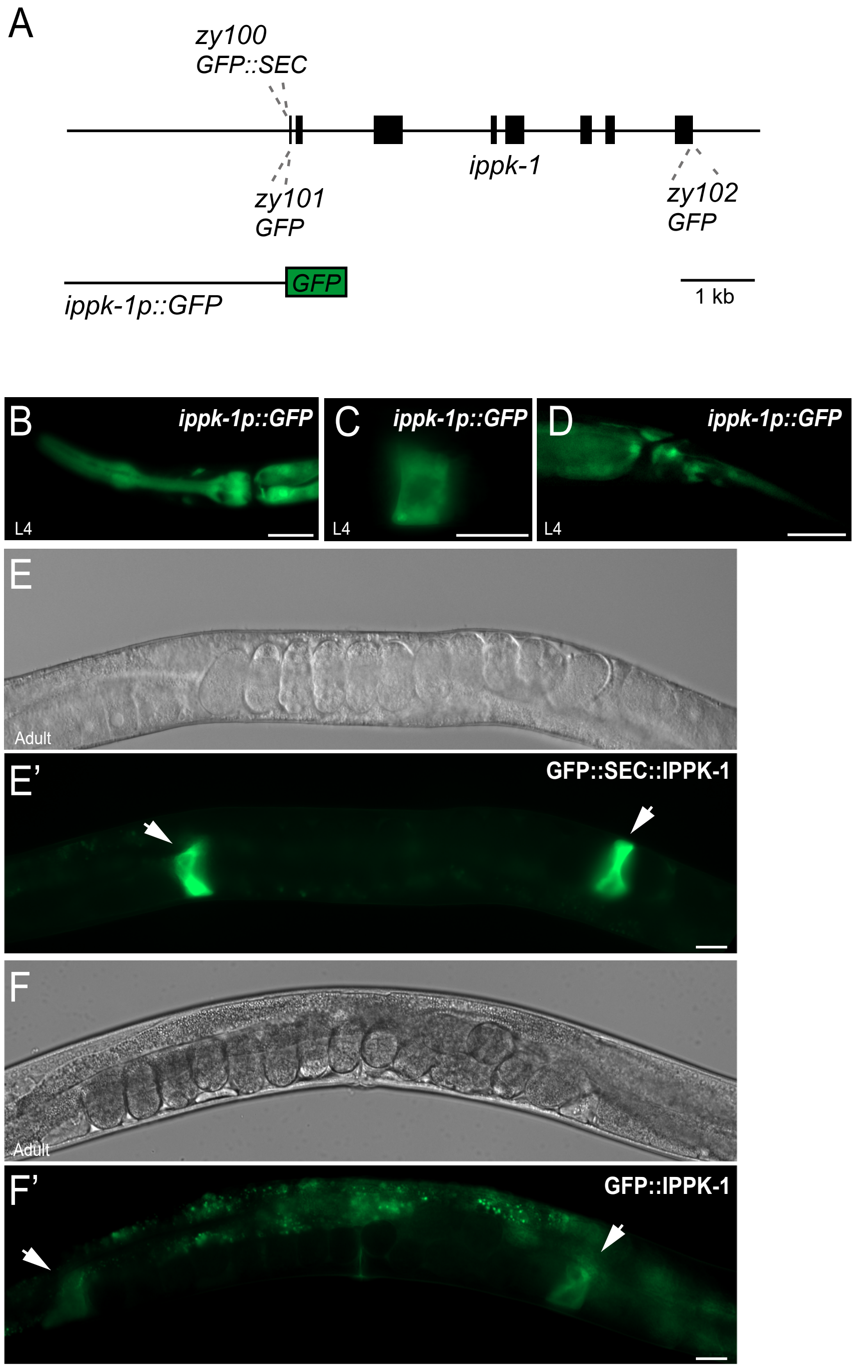
A
zy100
GFP::SEC
ippk-1
zy101
GFP
zy102
GFP
GFP
ippk-1p::GFP
1 kb
B
ippk-1p::GFP
L4
C
ippk-1p::GFP
L4
D
ippk-1p::GFP
L4
E
Adult
E'
GFP::SEC::IPPK-1
F
Adult
F'
GFP::IPPK-1

**FIG S4**

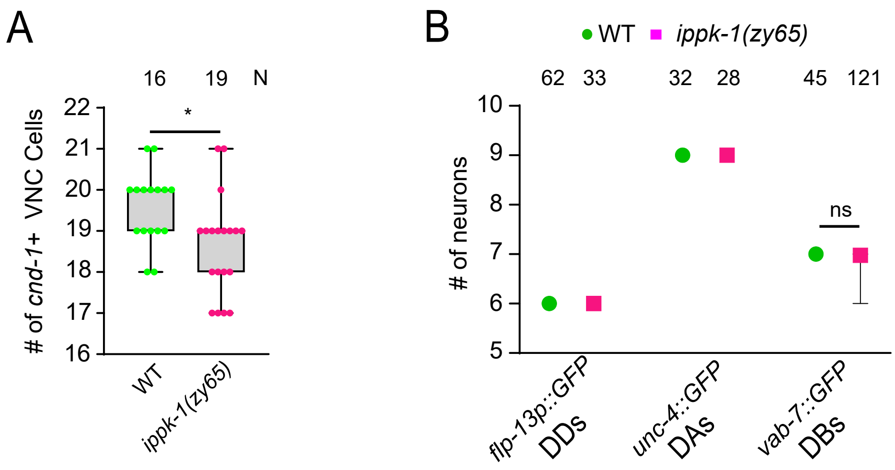

A
16
19
N
### of cnd-1+ VNC Cells
22
21
20
19
18
17
16
*
WT
ippk-1(zy65)
B
WT
ippk-1(zy65)
62
33
32
28
45
121
### of neurons
10
9
8
7
6
5
ns
flp-13p::GFP
DDs
unc-4::GFP
DAs
vab-7::GFP
DBs


FIG S5
