## Supplemental Table S1 for "IPPK-1 and IP6 contribute to ventral nerve cord assembly in *C. elegans*"

| Oligo # | DNA Oligos Sequence (5' – 3') | Source | Use |
|---|---|---|---|
| N1 | AAATGGCCAAATTAAATGCAGTTTTAGAGCTAGAAATAGCAAGT | Eurofins | sgRNA for *ippk-1::GFP* knock-in |
| N2 | CAAGACATCTCGCAATAGG | Eurofins | sgRNA universal reverse primer |
| N3 | ACGTTGTAAAACGACGGCCAGTCGCCGGCAGGTCATATTTCGGCG GATGT | Eurofins | 5' Repair template for *ippk-1::GFP* knock-in |
| N4 | CATCGATGCTCCTGAGGCTCCCGATGCTCCAATGCACGGCTTAAAA TTTCGC | Eurofins | 5' Repair template for *ippk-1::GFP* knock-in |
| N5 | CGTGATTACAAGGATGACGATGACAAGAGATAATTTGGCCATTTTT TGCATCAA | Eurofins | 3' Repair template for *ippk-1::GFP* knock-in |
| N6 | GGAAACAGCTATGACCATGTTATCGATTTCAGGAAATTGGAAATG TACGTGG | Eurofins | 3' Repair template for *ippk-1::GFP* knock-in |
| N7 | GAGGATTAATCTGCATTTTCGTTTTAGAGCTAGAAATAGCAAGT | Eurofins | sgRNA for *GFP::ippk-1* knock-in |
| N8 | ACGTTGTAAAACGACGGCCAGTCGCCGGCAGTTTCCGCAGAATCC CCATC | Eurofins | 5' Repair template for *GFP::ippk-1* knock-in |
| N9 | TCCAGTGAACAATTCTTCTCCTTTACTCATTTTCGGGTGGTTTTGGT GAG | Eurofins | 5' Repair template for *GFP::ippk-1* knock-in |
| N10 | CGTGATTACAAGGATGACGATGACAAGAGAATGCAGATTAATCCT CCTGCA | Eurofins | 3' Repair template for *GFP::ippk-1* knock-in |
| N11 | TCACACAGGAAACAGCTATGACCATGTTATCCACGAGGGGCATTT CAAAT | Eurofins | 3' Repair template for *GFP::ippk-1* knock-in |
| N12 | ACTATACAAAATGCAGATTAATCCTCCTG | Eurofins | GFP:IPPK-1 Fragment 1 |
| N13 | TCCATGTACCCTTTGGAATGTGTTCAGTG | Eurofins | GFP:IPPK-1 Fragment 2 |
| N14 | CATTCCAAAGGGTACATGGATCGGTTATTG | Eurofins | GFP:IPPK-1 Fragment 3 |
| N15 | GCCGACTAGTAAAATGCGATATACATTCATTATTG | Eurofins | GFP:IPPK-1 Fragment 4 |
| N16 | ATCGCATTTTACTAGTCGGCCGTACGGG | Eurofins | GFP:IPPK-1 Fragment 5 |
| N17 | TAATCTGCATTTTGTATAGTTCATCCATGCCATGTG | Eurofins | GFP:IPPK-1 Fragment 6 |
| N18 | ATAGCATGCAATACTAAATTAGCCAGAGGCG | Eurofins | Amplify the ippk-1 promoter |
| N19 | TTAGGATCCTTTCGGGTGGTTTTGGTGAGT | Eurofins | Amplify the ippk-1 promoter |
| N20 | ATAGCATGCTAGCAAGAAGCCAGCAAGAAG | Eurofins | Amplify the cnd-1 promoter |
| N21 | TTAGGATCCTTTCCAGTGATGTCCGCCAAC | Eurofins | Amplify the cnd-1 promoter |
| N22 | CATTTTGCTCTTGGAATAACGCA | Eurofins | Sequence *ippk-1(zy65)* |
| N23 | TTTGTCCAGGAGGCTTGTCT | Eurofins | Sequence *ippk-1(zy65)* |
| N24 | GTTGACGTACATCAGCTCGC | Eurofins | Sequence *ippk-1(or1572)* |
| N25 | TGGACACACAGAATAAGCGC | Eurofins | Sequence *ippk-1(or1572)* |
| N26 | TCTCCACCCAAAACTCAGGA | Eurofins | Genotype *ipmk-1(tm2687)* |
| N27 | CACCTCTGAACCTATACCGCT | Eurofins | Genotype ipmk-1(tm2687) |
| N28 | ACCCTGCAATATCCGGTTCA | Eurofins | Genotype *ipmk-1(tm2687)* |
| N29 | TTCACCTTGACAACGCCAGA | Eurofins | Genotype *vang-1(ok1142)* |
| N30 | GGCATGATGACTAGCCCACAA | Eurofins | Genotype *vang-1(ok1142)* |
| N31 | TGTGACAGGGAAAAGTGGAC | Eurofins | Genotype *vang-1(ok1142)* |
| N32 | TTCCTGTGATTGTCATTGTTGC | Eurofins | Genotype *sax-3(zy5)* |
| N33 | TCGATCGAGATCGTCTTCCA | Eurofins | Genotype *sax-3(zy5)* |
